# Acquisition of cross-resistance to Bedaquiline and Clofazimine following treatment for Tuberculosis in Pakistan

**DOI:** 10.1101/630012

**Authors:** Arash Ghodousi, Alamdar Hussain Rizvi, Aurangzaib Quadir Baloch, Abdul Ghafoor, Faisal Masood Khanzada, Mehmood Qadir, Emanuele Borroni, Alberto Trovato, Sabira Tahseen, Daniela Maria Cirillo

## Abstract

We report on the first six cases of acquired resistance to bedaquiline in Pakistan. Seventy sequential culture isolates from 30 drug-resistant tuberculosis patients on bedaquiline-containing regimen were tested for minimum inhibitory concentration (MIC) and whole genome sequencing. An increase in MICs associated with cross-resistance to clofazimine and appearance of specific mutations was documented in six cases. The study underlines that appropriate monitoring is mandatory for the introduction of new drugs.

## Manuscript

In 2018, following the availability of new evidence, the World Health Organization (WHO) updated its guidelines on the treatment of drug-resistant tuberculosis (TB) (1). The new guidelines recommend Bedaquiline as core drug in the standard combination regimen for the treatment of rifampicin-resistant TB. As a result, the number of patients eligible to receive bedaquiline-containing regimens would significantly increase. Encouraging results from studies currently underway on the use of bedaquiline in all-oral, shorter-course regimens for the treatment of rifampicin-resistant TB could result in an even greater use of bedaquiline in TB control programmes globally (2,3). However, solid surveillance of drug-resistance is needed to monitor the emergence of resistance to bedaquiline in patients during therapy.

We report first six cases of acquired bedaquiline resistance identified in Pakistan as the results of a surveillance project to monitor acquisition of resistance to bedaquiline implemented by the Pakistan National TB Reference Laboratory (NRL) and TB Supranational Reference Laboratory (SRL-Milan). In Pakistan, bedaquiline was introduced in November 2015 for the treatment of MDR-TB. Culture is performed routinely before start and during the course of MDR treatment. Arrangement were made to collect clinical isolates from culture laboratories serving MDR patient on bedaquiline-containing regimen enrolled at six sites located in six different cities. Clinical isolates were collected at NRL and sub-cultured to obtain fresh isolates. Sequential isolates from individual patient including a baseline and follow up culture isolated at 3 or 4 month were targeted for testing. Six-and nine-month isolates were also included if culture-positive. Cultures isolated earlier during treatment were included only if three-months culture was not available or showed resistance. Selected isolates were transported to the SRL in batches between November 2017 and May 2018. Patients’ clinical history was retrospectively extracted from medical records. Institutional review board for HIV, TB and Malaria programme, Pakistan approved the study.

At SRL-Milan, all strains were tested for Minimum Inhibitory Concentration (MIC) for bedaquiline in 7H11 medium and cases with elevated bedaquiline MIC in isolates collected during therapy compared to baseline, were tested for bedaquiline and clofazimine MIC in BACTEC MGIT960 (BD, Franklin Lakes, NJ, USA) (4). Bedaquiline dry powder was supplied by Janssen-Pharmaceutica (Beerse, Belgium).

Whole genome sequencing (WGS) was carried out with the Illumina Nextera-XT DNA sample preparation kit to prepare paired-end libraries of 150-bp read-length for Illumina sequencing. Data analysis and SNPs calling were performed using the MTBseq-Pipeline on low-frequency detection mode (5). Genes associated to resistance to bedaquiline and/or clofazimine (*atpE, Rv0678, pepQ, Rv1979c)* were screened for mutations (6-9).

A total of 158 unique clinical isolates were selected for testing at SRL-Milan including 70 sequential isolates collected from 30 individual patients (8MDR, 15 pre-XDR and 7 XDR). Sequential isolates tested included two each from twenty two, three from six and four from two patients. Acquired drug resistance was reported in post exposure MTB isolates from six patients showing an increase in bedaquiline MICs in 7H11 medium (0.25–0.5 mg·L^−1^). The phenotypic/genotypic characteristics of MTB isolates from these six HIV negative, unrelated, pulmonary TB patients are summarised in table1.

Patient-1, a newly diagnosed rifampicin resistant-TB case, was enrolled on second-line treatment. Patient remained culture positive and was subsequently diagnosed as pre-XDRTB case and after five months bedaquiline, clofazimine and linezolid was added to the regimen. The patient continued to be culture positive and six months later, delamanid was also added to the regimen and ultimately treatment failure was declared. The first increase in bedaquiline MIC was seen at fourth month of treatment (0, 5 mg·L^−1^ in 7H11) and at this point WGS analysis showed an insertion (Ins) at nucleotide position 141-142 of *Rv0678* (after genomic position 779130). At month 9 isolate became resistant to delamanid (MIC >1 mg·L^−1^, data not shown) and WGS confirmed the presence of the *Rv0678* mutation and also showed a mutation in *fgd1* (G104S) probably responsible for delamanid resistance.

Patients 2 and 3 had MDR-TB with history of failed second line treatment taken for 20 and 17 months, both were then re-enrolled on bedaquiline and clofazimine containing regimen. For both the patients, no difference was seen in bedaquiline MIC of the isolate collected before and after two and one month of bedaquiline containing treatment respectively. Subsequent isolate from patient-2 at fifth month of treatment became resistant to both bedaquiline and clofazimine and WGS analysis showed an insertion at position 141-142 of *Rv0678*. Similarly, isolate from patient-3 after 6 months of bedaquiline, showed eight and four-fold increase in bedaquiline and clofazimine MICs in MGIT and a V20G mutation in *Rv0678* was detected. Moreover, the MTB strains from patient-2 at baseline and at two and five months of treatment carried P478G mutation in *Rv1979c* suggesting that this mutation has no effect on clofazimine resistance. Patients 4, 5 and 6 were XDR-TB cases failing second line treatment regimen containing clofazimine taken for 20, 14 and 12months respectively. Baseline isolates of all three cases had bedaquiline MIC of 0.03 mg·L^−1^ in 7H11 and showed no detectable mutations in *Rv0678, pepQ, atpE* and *Rv1979c*. The follow-up isolates, from patients 4 and 5 collected at months 6 and 5 during treatment showed an increase in bedaquiline and clofazimine MIC due to an insertion in at position 138-139 of *Rv0678* (after genomic position 779127). Patient-6, showed a substantial increase in MICs for both bedaquiline and clofazimine (8 fold bedaquiline, 4 clofazimine) after one month of therapy and we detected an insertion in *Rv0678* at low allele frequency (12.7%) which was not present at baseline. This patient ultimately failed treatment.

Breakpoints for bedaquiline are still provisional (10,11) and patients are started on treatment without susceptibility tests. We adopted 0.25 mg·L^−1^ as susceptibility breakpoint in 7H10 and 7H11(12, 13), which is above the MIC that inhibits 90% of the isolates or strains (0.12 mg·L^−1^). However, with this breakpoint, the strains collected from two patients reported as “failure” at the end of treatment (patient-4 MIC 0.25 mg·L^−1^, eight fold higher than baseline and patient-5 MIC 0.125 mg·L^−1^, four fold higher than baseline) would have been categorised as susceptible. This finding indicates that monitoring MICs during treatment could be a better predictor for failure than single testing at critical concentration. The genetic basis of resistance to bedaquiline is still the subject of much uncertainty. WGS analysis in different studies showed that bedaquiline-clofazimine cross resistance arises through mutations in *Rv0678* (6,7,8) *and pepQ* (9). In this study, we show that the increase of MIC to bedaquiline and clofazimine could be explained by mutations emerging during therapy in *Rv0678*. As reported in table 1 we observed four different mutations and three of these (138-139_Ins-G, 141-142_Ins-C, 192-193_Ins-G) were previously reported as associated to bedaquiline resistance in MTB clinical strains (8). We show that the same mutations are associated to Clofazimine resistance. As a result, regimens that contain both drugs might have to be reconsidered when these mutations are identified, to reduce the risk of failure for the patients and transmission of such strains in the community. Altogether, these data show that resistance to bedaquiline emerges during treatment and emphasize the importance of using MIC and whenever possible molecular based surveillance in national programmes implementing bedaquiline for the treatment of MDR/XDR-TB to monitor the emergence of resistance. Moreover, a classification of mutations associated to bedaquiline and clofazimine resistance is crucial to lead future development of tools for fast detection of resistance. Introducing drugs without proper diagnostics to monitor drug resistance may lead to amplification of hardly treatable cases.

**Table 1:** The phenotypic/genotypic characteristics of MTB isolates from these six patients who acquired cross-resistance to bedaquilline and clofazimine

| Strain -ID | Previous Second line treatment Regimen | DST profile at start of BDQ containing Regimen | BDQ exposure | M. tuberculosis strain- Testing results |  |  |  |  |  |  |  |  | Treatment outcome |
| --- | --- | --- | --- | --- | --- | --- | --- | --- | --- | --- | --- | --- | --- |
|  | /Outcome |  | (month) | pDST results |  |  | WGS |  |  |  |  |  |  |
|  |  |  |  | BDQ-MIC in 7H11 (mg·L<sup>-1</sup>) | BDQ-MIC in MGIT( mg·L<sup>-1</sup>) | CFZ-MIC in MGIT( mg·L<sup>-1</sup>) | Genome coverage | Rv0678 | pepQ | atpE | Rv1979c | Lineage |  |
| H37Rv (NC_000962.3) |  |  |  | 0.03 | 0.25 | 0.25 | 43.28 | WT | WT | WT | WT |  |  |
| Patient-1 | NO | Pre-XDR | 0 | 0.125 | 0.5 | 0.5 | 71.67 | WT | WT | WT | WT | Delhi-CAS(3) | Failure |
|  |  |  | 4 | 0.5 | Not tested | Not tested | 40.57 | 779130_141-142_Ins-C | WT | WT | WT |  |  |
|  |  |  | 6 | 0.5 | Not tested | Not tested | 58.11 |  | WT | WT | WT |  |  |
|  |  |  | 9 | >0.5 | 4 | 4 | 68.20 |  | WT | WT | WT |  |  |
| Patient-2 | Am(8),LFX,Cs,Eto,Z,E,B6 /Failure | MDR | 0 | 0.03 | 0.5 | 0.5 | 80.97 | WT | WT | WT | 2221732_P478G | Delhi-CAS(3) | Cured |
|  |  |  | 2 | 0.06 | Not tested | Not tested | 127.50 | WT | WT | WT |  |  |  |
|  |  |  | 5 | 0.5 | 4 | 4 | 90.23 | 779130_141-142_Ins-C | WT | WT |  |  |  |
| Patient-3 | Am,Lfx,Eto,CS,Z,PAS,B6 /Failure | MDR | 0 | 0.06 | 0.25 | 0.5 | 96.26 | WT | WT | WT | WT | Delhi-CAS (3.1.2) | Failure |
|  |  |  | 1 | 0.06 | Not tested | Not tested | 86.92 | WT | WT | WT | WT |  |  |
|  |  |  | 6 | 0.06 | 2 | 2 | 100.78 | 779048_V20G | WT | WT | WT |  |  |
| Patient-4 | Cm(12),MfX,Cs,Eto,Cfz,Z, Amx/clv /Failure | XDR | 0 | 0.03 | 0.25 | 0.5 | 81.93 | WT | WT | WT | WT | Delhi-CAS(3) | Still on treatment |
|  |  |  | 2 | 0.03 | Not tested | Not tested | 111 | WT | WT | WT | WT |  |  |
|  |  |  | 7 | 0.25 | 2 | 4 | 102.74 | 779127_138-139_Ins-G | WT | WT | WT |  |  |
| Patient-5 | Cm(12),Mfx,Cs,Eto,Cfz,Z,PAS,Clr,Amx/clv,H,E,B6 /Failure | XDR | 0 | 0.03 | 0.5 | 0.5 | 68.40 | WT | WT | WT | WT | Delhi-CAS(3) | Failure |
|  |  |  | 1 | 0.06 | 1 | Not tested | 125.22 | WT | WT | WT | WT |  |  |
|  |  |  | 3 | 0.125 | 2 | Not tested | 116.61 | 779127_138-139_Ins-G | WT | WT | WT |  |  |
|  |  |  | 6 | 0.125 | 2 | 4 | 109.03 |  | WT | WT | WT |  |  |
| Patient-6 | Cm(12),MfX,Cs,Eto,Cfz,LNZ,PAS,Z, Amx/clv,B6 /Failure | XDR | 0 | 0.03 | 0.5 | 1 | 99.35 | WT | WT | WT | WT | Euro-American -4.9 | Failure |
|  |  |  | 1 | 0.5 | 4 | 4 | 83.34 | 779181_192-193_Ins-G | WT | WT | WT |  |  |

**Figure 1.**
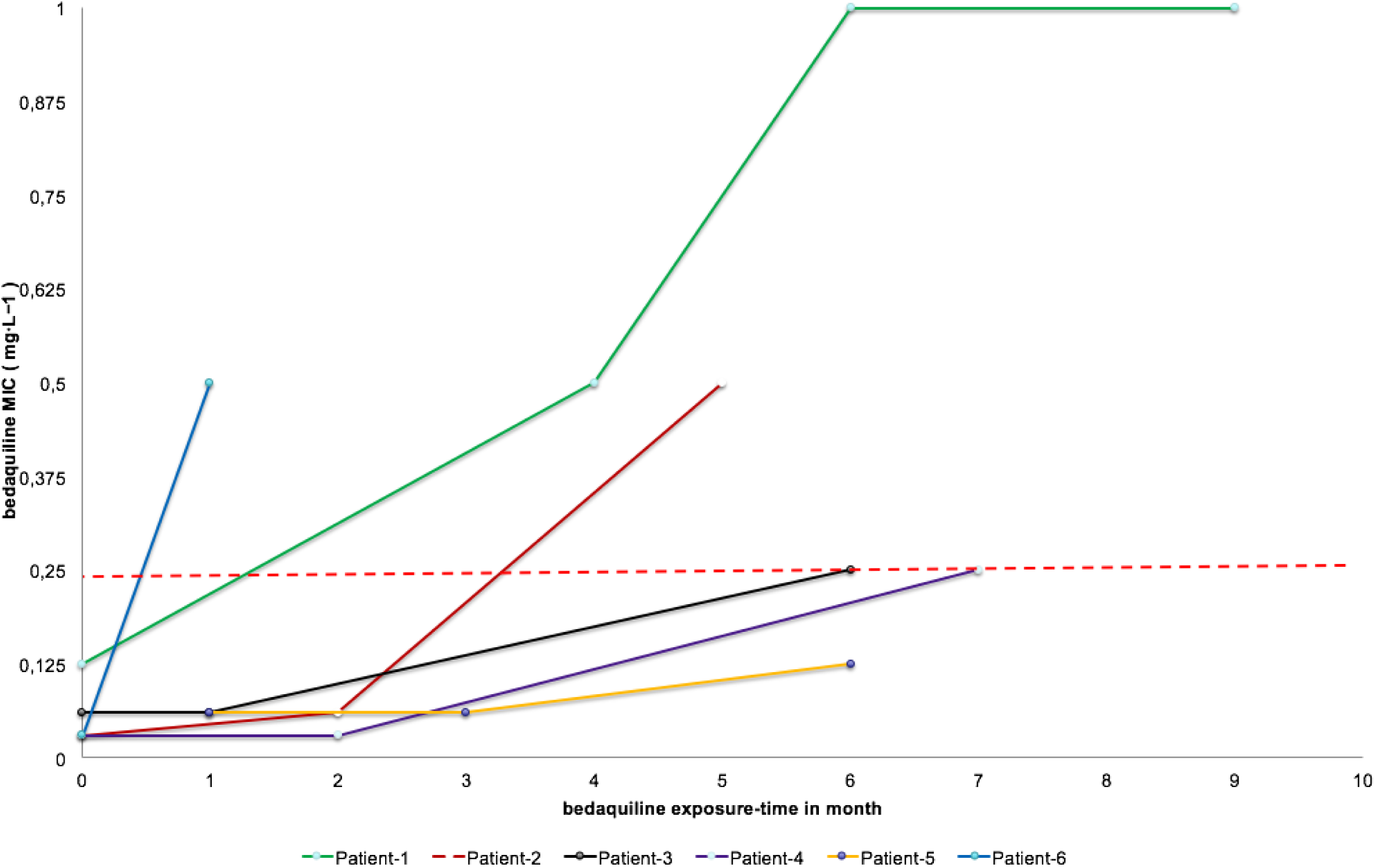
The MIC of bedaquiline performed in 7H11 medium in patients under treatment with bedaquiline-contain regimen at different time points. The dashed line in red shows the currently identified critical concentration for bedaquiline in 7H11 medium by WHO (13).

## Ethics approval

Ethical clearance was approved by the Institutional review board for HIV, TB and Malaria programme, Pakistan.

## Conflicts of interest

None declared

## Acknowledgements

We would like to acknowledge Andrea Maurizio Cabibbe for his guidance, Anna Dean and Matteo Zignol for their critical revision during the preparation of this manuscript.

## Support statement

This study was funded by a WHO project to NRL/NTP Pakistan and SRL-Milano on the surveillance of acquisition of resistance to bedaquiline in Pakistan

